# Single-cell chromatin accessibility analysis of mammary gland development reveals cell state transcriptional regulators and cellular lineage relationships

**DOI:** 10.1101/624957

**Authors:** Chi-Yeh Chung, Zhibo Ma, Christopher Dravis, Sebastian Preissl, Olivier Poirion, Gidsela Luna, Xiaomeng Hou, Rajshekhar R. Giraddi, Bing Ren, Geoffrey M. Wahl

## Abstract

It has only recently become possible to obtain single-cell level resolution of the epigenetic changes that occur during organ development. We reasoned that precision single-cell chromatin accessibility mapping of mammary gland development could provide needed insight into the epigenetic reprogramming and transcriptional regulators involved in normal mammary gland development. Here, we provide the first single-cell resource of chromatin accessibility for murine mammary development from the peak of fetal mammary stem cell (fMaSC) functional activity in late embryogenesis to the differentiation of adult basal and luminal cells. We find that the chromatin landscape within individual cells predicts both gene accessibility and transcription factor activity, and we present a web application as a scientific resource for facilitating future analyses. Strikingly, these single-cell chromatin profiling data reveal that fMaSCs can be separated into basal-like and luminal-like lineages, providing evidence of early lineage segregation prior to birth. Such distinctions were not evident in analyses of single-cell transcriptomic data.

## Introduction

The specialized functions of tissues require the coordinated activities of diverse differentiated cell types derived from stem/progenitor antecedents (Donati and Watt, 2015). The “epigenetic programming” of stem cells enables them to either retain their multi-potentiality or differentiate into the specific cell types. In some cases, “epigenetic reprogramming” allows cells to gain developmental plasticity to repair tissue injury (Ge and Fuchs, 2018). Determining the epigenetic and molecular programs that generate unique cell identities or developmental plasticity are critical for understanding the mechanisms for generating cell type heterogeneity during normal tissue homeostasis, and for enabling repair after injury. Perturbation of these mechanisms by oncogene activation, tumor suppressor loss, and inflammatory stimuli likely contribute to the cell state reprogramming increasingly observed during progression of many cancers (Feinberg et al., 2016; Kawamura et al., 2009; Koren et al., 2015; Schwitalla et al., 2013; Van Keymeulen et al., 2015).

The mammary gland is an excellent system for studying mechanisms of cellular specification due to its accessibility, the dramatic changes it undergoes in embryogenesis and post-natal development in response to puberty, pregnancy, and involution, and the substantial knowledge gained about factors involved in these cell state transitions (Inman et al., 2015; Makarem et al., 2013; Veltmaat et al., 2003). However, there is also considerable debate on the nature of the mammary stem cells that generate and sustain the gland, and on the mechanisms for establishing the basal and luminal cell lineages (Visvader and Stingl, 2014). One model proposes that bipotent mammary stem cells arise during embryogenesis (herein referred to as fetal mammary stem cells, fMaSCs), and they generate basal, luminal progenitor (LP) and mature luminal (ML) populations that are postnatally maintained by lineage restricted progenitors (Davis et al., 2016; Giraddi et al., 2015; Van Keymeulen et al., 2011; Wuidart et al., 2016). But the precise time and mechanisms by which fMaSC bipotency becomes luminally- or basally-restricted remains unknown. Based on recent lineage tracing studies, it has been proposed that basal and luminal lineage specifications occur before birth (Elias et al., 2017; Lilja et al., 2018; Wuidart et al., 2018) but direct evidence for the presence of lineage restricted populations during embryogenesis supported by epigenetic and molecular profiling of these populations has not been presented.

One way of determining when primitive, undifferentiated embryonic cells become lineage committed is to use agnostic single-cell molecular profiling. Analysis of large cell populations isolated from different developmental stages using single-cell RNA sequencing (scRNA-seq) combined with bioinformatic analyses to generate lineage relationships and pseudotime developmental trajectories has been used for this purpose. One recent scRNA seq study that analyzed hundreds of E18 mammary cells, which have the highest *in vitro* and *in vivo* fMaSC activity, showed that they comprise a single diffuse transcriptomic cluster, with most of the cells sharing characteristics of both basal and luminal cells, as might be expected of undifferentiated bipotent cells (Giraddi et al., 2018). An independent study using a limited number of E14 cells for RNA sequencing came to a similar conclusion about the mixed-lineage nature of the bipotent cells, and further showed that the E14 cells could be traced into adult luminal and basal cells (Wuidart et al., 2018). Pseudotime analyses produced a trajectory in which the E18 “cluster” generated a basal subset and a luminal progenitor subset shortly after birth. The luminal progenitor was then inferred to generate a mature luminal component when analyzed in the prepubertal adult (Giraddi et al., 2018). This study was consistent with an independent analysis that focused on post-natal and adult cells (Bach et al., 2017), but differed from those of another study (Pal et al., 2017) that concluded that a uniform, basally oriented cell cluster was present after birth, and that this basal cluster generated the luminal lineages. However, the latter results are not consistent with the luminal-specific lineage tracing studies that show the lack of basal contributions to the luminal cell populations postnatally (Elias et al., 2017; Lilja et al., 2018; Van Keymeulen et al., 2011; Wuidart et al., 2016, 2018).

The inability of single-cell RNA-seq to identify putative uni-potent cells is not an argument against their existence, but may rather reflect technical or computational limitations such as low numbers of reads per cell and the potential for regulatory factors expressed at low transcript numbers to fall below the detection threshold (“drop-outs”). We therefore sought another strategy to bypass these limitations. We performed single-nucleus ATAC-seq (snATAC-seq) for efficient and high-quality profiling of chromatin accessibility at single-cell resolution. Because chromatin accessibility is less transient than transcript expression, and it indicates regulatory potential rather than the current cell state as implicated by the transcriptome, we reasoned that analyzing chromatin state profiles may better separate cell types than gene expression alone (Corces et al., 2016; Shema et al., 2019). We therefore wanted to determine whether snATAC-seq enables greater resolution of cell state heterogeneity in mammary tissues. The large sample size of single-cell studies also allows for the application of machine learning techniques that can identify cell-type specific genes and regulatory mechanisms that are involved in establishing mammary cell state heterogeneity (Pott and Lieb, 2015). Here we agnostically profile mammary cells at different developmental stages using snATAC-seq to determine whether this approach clarifies the timing, extent, and underlying molecular mechanisms associated with lineage specification in mammary development. The data and strategies described provide a resource for future epigenetic studies of mammary cell regulation, a catalog of upstream control elements containing binding sites for cell state-determining transcription factors, computational approaches that provide finer distinction of mammary cell states, and a new pseudotime progression of mammary differentiation.

## Results

### Profiling the chromatin accessibility landscape at single-cell resolution during mammary development

We used a combinatorial indexing assisted single-cell ATAC-seq strategy to interrogate chromatin accessibility in mammary tissue at single-cell resolution spanning the interval from late embryogenesis to the adult (single-nucleus ATAC-seq; snATAC-seq) (Figure 1A) (Cusanovich et al., 2015; Preissl et al., 2018). This approach allows scalable profiling of thousands of single nuclei while maintaining sufficient read depth to obtain high quality chromatin accessibility data. All cell populations were first purified using FACS to obtain EpCAM+, Lin-cells, which enabled removal of non-epithelial stromal and blood cells (Figure S1A). The aggregated fetal and adult single-nucleus profiles revealed nucleosomal fragmentation patterns, high correlation between the biological replicates, and good signal-to-noise levels (Figure S1B-D and Table S1). Importantly, the snATAC-seq data showed enrichment of accessible regions that were previously identified by bulk ATAC-seq profiling of FACS-enriched fetal and adult cells (Dravis et al., 2018) (Figure S1E).

**Figure 1.**
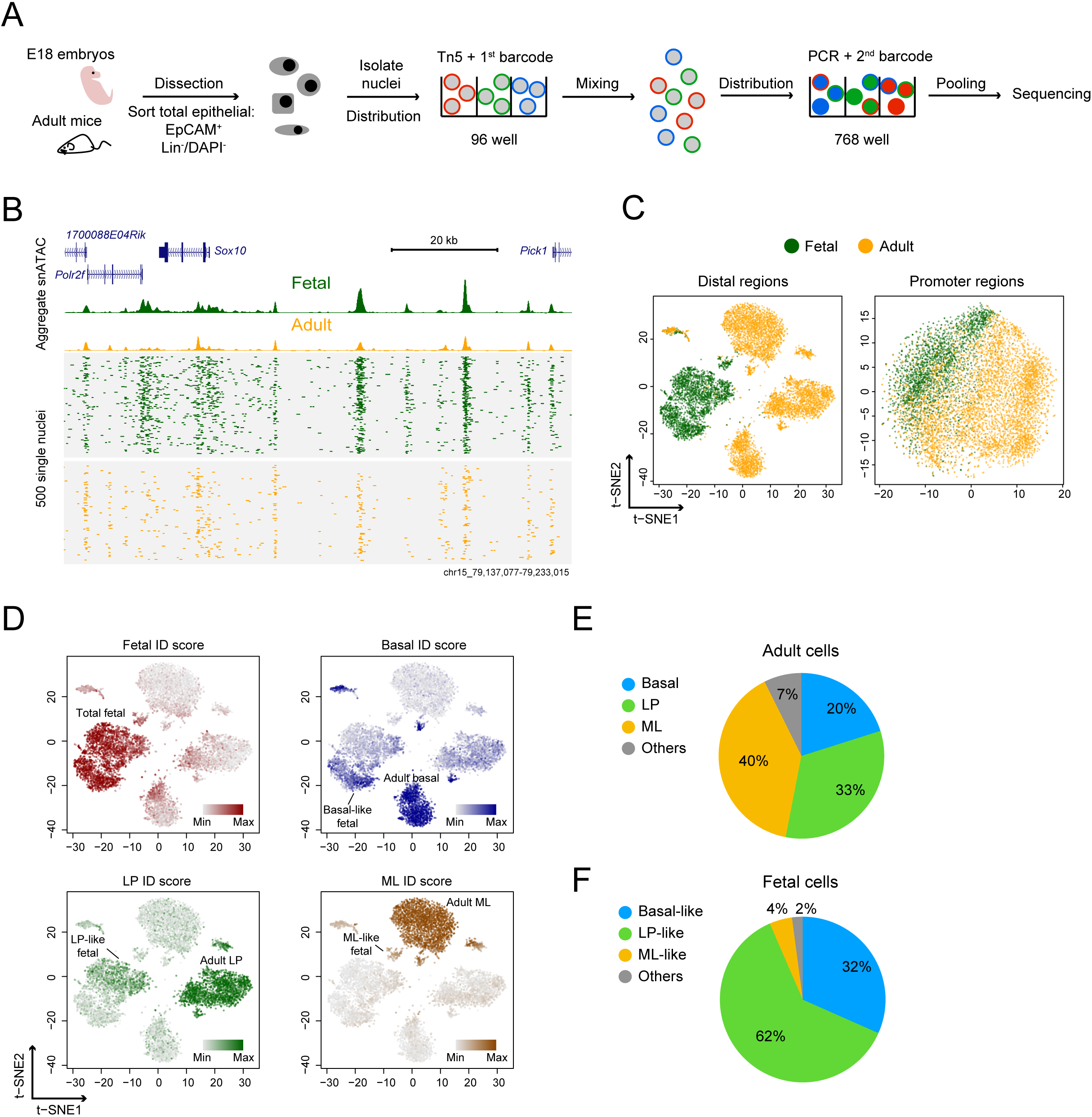
snATAC-seq in fetal and adult mammary epithelial cells. (A) Overview of snATAC-seq experimental strategy. (B) Aggregate (top) and single-nucleus (bottom) ATAC-seq profile of mammary cells. Reads from 500 randomly selected cells are plotted to represent the single-nucleus profile. (C) t-SNE representation of snATAC-seq profile generated from either distal (> ±1.5 kb transcription start site [TSS]) or promoter regions (< ±1.5 kb TSS). Shown plots are combined from two biological replicates. (D) Single-cell ID score of major mammary cell types overlaid on the snATAC-seq profile. Cell cluster annotations are shown. (E) Cellular composition of adult cell population derived from snATAC-seq data. (F) Cellular composition of adult-like cell type in the fetal cell population.

We employed a computational framework that has previously been shown to be able to identify cell clusters in diverse tissues to analyze snATAC-seq data (Cusanovich et al., 2015). We first filtered out low quality nuclei with stringent selection criteria including read depth per cell (> 2000) and percentage of reads in peaks (> 20%) (Figure S1F). This resulted in 7,846 high quality single nuclei derived from 2,577 fetal and 5,269 adult cells (Figure 1B). For all replicates, the median reads per nucleus ranged from 4,420 to 7,488, and median reads in peaks ranged from 69% to 74%. We then used the aggregate snATAC-seq profile from each replicate to determine open chromatin regions. This revealed 21,179 promoter-proximal regions (< ±1.5 kb transcriptional start site [TSS]) and 145,453 distal regions (>= ±1.5 kb TSS). Within both promoter-proximal and -distal regions, we constructed a single-cell binary matrix of chromatin accessibility and then performed latent semantic indexing (LSI) and dimension reduction for signal normalization and processing (Figure S2A) (see Methods). We used t-distributed stochastic neighbor embedding (t-SNE) to ascertain patterns of relatedness of snATAC-seq data derived from distal regions (Figure 1C) (Maaten and Hinton, 2008). The two biological replicates analyzed reproducibly resulted in one fetal and three adult major cell clusters (Figure S2B). Such cluster structures were not observed using only promoter-proximal snATAC-seq data or using control ‘shuffled’ nuclei profiles (Figure 1C and S2C). These results show that the clusters observed are highly specific, and are consistent with previous reports showing that accessibility at distal elements is more specific to cell type than is accessibility at promoter-proximal regions (Shlyueva et al., 2014; Wu et al., 2016).

### Identification of major mammary cell types from snATAC-seq profile

Density peak clustering on cells after t-SNE identified 14 clusters in all, some of which represent sub-clusters within major cell types (Figure S3A, B). We excluded clusters 10 and 14 as their unusually high read count distributions suggests they may result from barcode collision events (Figure S3C). To annotate the remaining 12 cell clusters, we examined the chromatin accessibility at cell-type specific open and closed regions we previously identified and verified within specific mammary cell populations using bulk ATAC-seq data (Dravis et al., 2018). We calculated a single-cell “ID score” for fetal, adult basal, LP, and ML cells. Higher ID scores represent greater similarity to the reference cell type (see Methods). We determined which cell clusters correspond to fetal, adult basal, LP and ML populations by overlaying the four ID scores onto the t-SNE (Figure 1D and S3D). We inferred from this approach that adult basal, LP and ML cells represent 20%, 33% and 40% of the total adult mammary population, respectively (Figure 1E), which is consistent with previous FACS-based studies (Shackleton et al., 2006; Stingl et al., 2006).

Interestingly, and in contrast to other cell types that possess relatively tight cluster structures, the approach described above separated fetal cells into three subclusters (two large and one small, Figure 1D). Although each of the three clusters possesses fetal characteristics, they can be distinguished by separate enrichment with basal, LP, and ML ID scores. The basal-, LP- and ML-like fetal cells represent 32%, 62% and 4% of the total fetal population, respectively (Figure 1F). This separation of fetal cells into adult-like sub-clusters is also evident when the data are visualized using uniform manifold approximation and projection (UMAP) (Figure S4) (McInnes et al., 2018), another dimensionality reduction method that better preserves lineage pairwise distances of embedding than t-SNE (Becht et al., 2018). Together, these results show that the snATAC-seq data is of high quality, that it is able to reveal cell types based on chromatin accessibility, and that it provides finer distinctions among fetal cells than previously possible using single-cell RNA-seq.

### E18 fetal mammary cells show features of partial lineage specification

Our use of cell ID score (Figure 1D) indicates that fetal cells can be subdivided into distinct clusters with some adult-like features. In order to validate this observation and investigate lineage specification, we calculated a ‘basal-to-luminal score’ by combinatorial analysis of the top basal and luminal accessible genes (see Methods). As expected, our analysis shows strong enrichment of a basal score in basal cells and a luminal score in ML (Figure 2A). LP cells are mostly located at the intermediate-to-luminal status, suggesting that these are luminal cells that possess some basal characteristics, which is consistent with our previous single-cell RNA-seq analyses (Giraddi et al., 2018). Interestingly, the three subclusters of fetal cells show clear indications of acquiring basal-, LP- or ML-like gene accessibility (Figure 2A).

**Figure 2.**
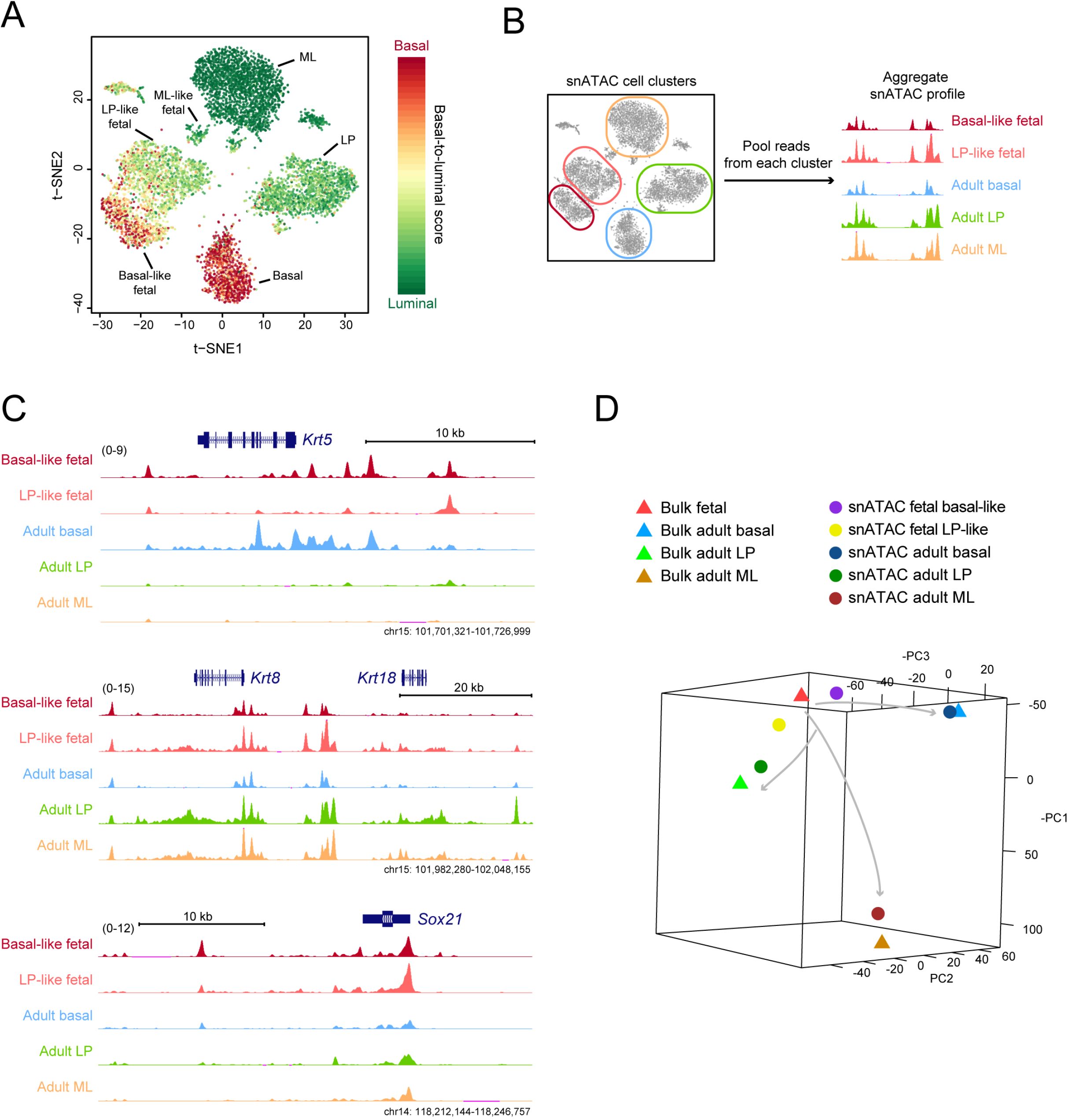
Fetal mammary cells at E18 show epigenetic features of partial lineage specification. (A) Single-cell basal-to-luminal score overlaid on the snATAC t-SNE plot. Cell cluster identities are shown. (B) Approach to generate aggregate snATAC profile based on single-cell clustering. (C) Signal tracks of aggregate snATAC profile. The signal ranges are shown in the parentheses. (D) PCA comparing aggregate snATAC profile with bulk ATAC-seq of sorted mammary populations. Arrows indicate putative mammary differentiation paths.

We also separated reads based on the single-cell clustering and aggregated them together to compare the cell cluster profiles as aggregated bulk (ML-like fetal cells were not analyzed as the cell number is too low to obtain an accurate aggregate signal; Figure 2B). We found that the basal-like fetal cells are more accessible at basal markers, including *Krt5* and *Acta2,* while LP-like fetal cells are more accessible at LP/pan-luminal markers, such as *Krt8, Krt18* and *Kit* (Figure 2C and S5). Both fetal sub-clusters are accessible at fetal-preferenced genes such as *Sox21* and *Sox10.* Comparing our aggregate snATAC-seq profile with published bulk ATAC-seq data using principle component analysis (PCA), we find that LP- and basal-like fetal cells have indeed moved toward the adult LP and basal chromatin state, respectively, but they still remain close to bulk fetal cells (Figure 2D). This suggests that fetal cells at this stage of mammary development are starting to acquire adult-like chromatin accessibility, but they still largely possess their fetal-specific features. These results support the conclusion that the epigenetic landscape of fetal cells at E18 is partially specified into states similar to the three major adult cell types. This type of poised landscape would position the cells to differentiate rapidly into the corresponding cell types after birth, as has been observed using RNA-sequencing and immune fluorescence analyses (Giraddi et al., 2018).

### Single-cell transcription factor dynamics of mammary development

Transcription factors and the programs they control mediate many cell specification events. We therefore wanted to use the snATAC-seq data to infer potential transcriptional regulators of cell state control during mammary development. We used chromVar, a package designed for analyzing sparse snATAC-seq data by inferring TF activity using variability of TF DNA-binding motif enrichment at accessible chromatin regions (Schep et al., 2017). ChromVar calculates a TF z-score, which infers for each single cell the TFs that are binding to open chromatin regions based on the TF motifs present within these regions. Our analysis reveals that TFs previously known to function in regulating mammary cell state are highly enriched in their corresponding cell clusters. For example, NFIX and SOX10 are enriched in fetal cells, while ELF5, FOXA1 and P63 are enriched in LP, ML and basal cells, respectively (Figure 3A). The TF z-scores of these major factors also highly correlate with their mRNA expression (Figure 3B), suggesting their relevance as transcriptional regulators. In addition, TF z-scores for members of the SMARCC, FOS-JUN, FOX, P63, NF1, NFkB and SOX-family TFs are significantly variable between the single mammary cells when compared to permutated background (Figure S6A and Table S2), suggesting that these factors may contribute to cellular differentiation programs.

**Figure 3.**
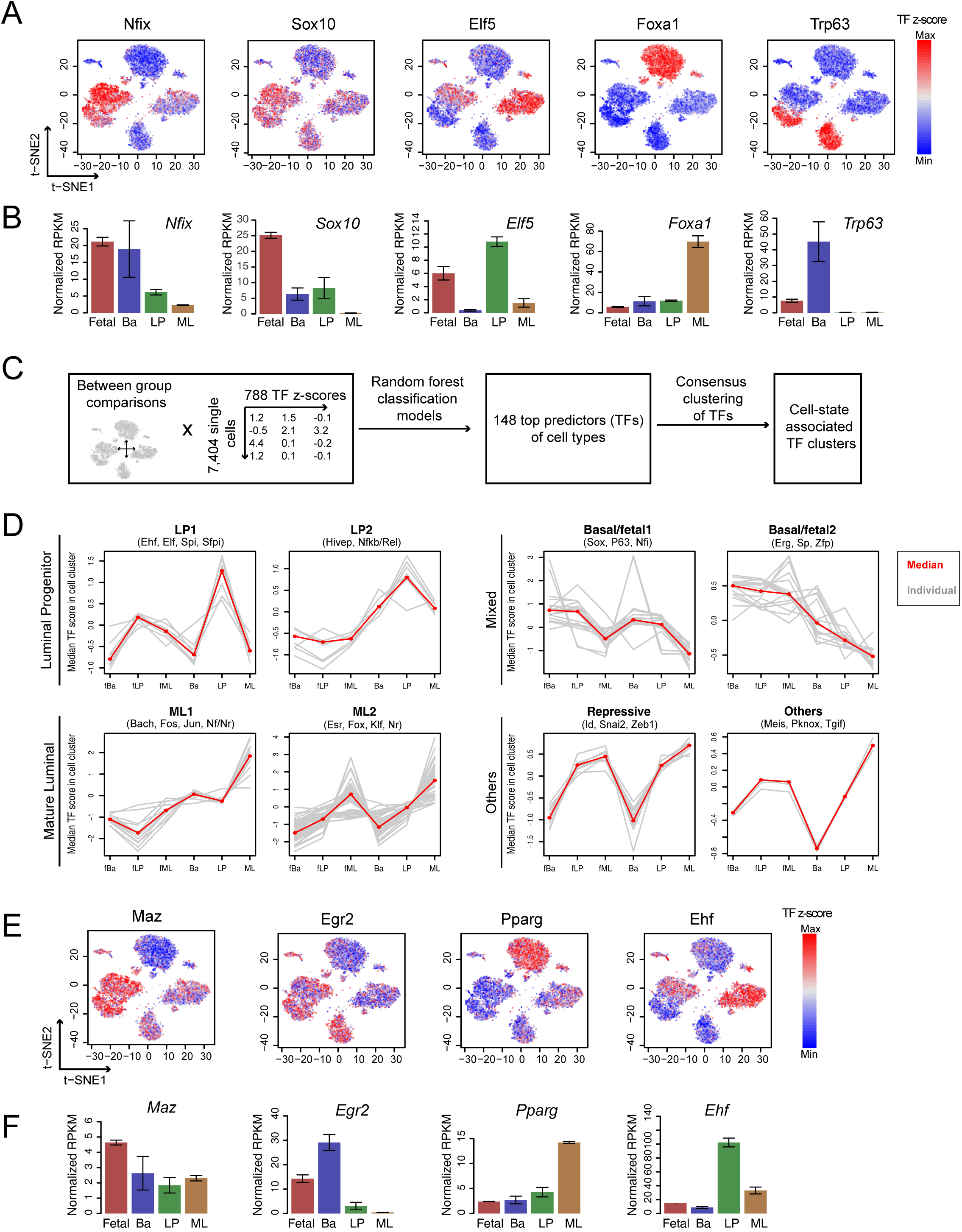
Single-cell transcription factor dynamics during mammary development. (A) TF z-score calculated from chromVar overlaid on the snATAC t-SNE plot. (B) RNA-seq expression of TFs from (A). Mean ± SEM (n = 2). (C) Computational framework to identify cell state predictive TFs. (D) TF z-score profile in cell types for the 8 TF clusters, grouped into luminal progenitor, mature luminal, mixed basal/fetal, and others type TFs. Individual (grey) and median (red) TF z-scores in cell type are shown. Examples of TF family in cluster are shown on the top. (E-F) TF z-score (E) and RNA-seq expression (F) of identified mammary cell state factors that are less known. Mean ± SEM (n = 2).

The above cell ID score and aggregate snATAC-seq analyses indicate that E18 fetal cells can already be separated into clusters with basal-, LP- and ML-like features. Supporting this observation, orthogonal analysis from the TF z-score also shows that each of the three fetal cell sub-clusters is enriched with adult LP (ELF5), basal (P63) or ML (FOXA1) associated TFs, despite all of these cells retaining significant fetal-like representation of embryonic factors such as NFIX (Figure 3A).

To systematically identify TFs that are associated with each mammary cell state, we employed a bioinformatic framework that leverages the large sample size of single-cell datasets by combining machine learning and data clustering (see methods) (Figure 3C). We used random forest, a high accuracy and low predictor bias machine learning approach (Geurts et al., 2009), to extract TFs that are important in predicting whether a cell belongs to the fetal, basal, LP or ML cluster. To improve the accuracy of the classification, cells corresponding to the stromal/mesenchymal cluster (cluster 6, figure S3B) or that likely result from barcode collisions (cluster 10 and 14, figure S3B) were removed before applying random forest. The random forest model was tuned to ensure optimal accuracy (Figure S6B-D). This resulted in 148 TFs as strong predictors of cell state (Table S3). We then performed consensus clustering on these TFs based on their TF z-scores across all the single cells. Using unsupervised cluster number selection, we identified 8 major TF clusters (Figure S6E and S6F) that can largely be separated into five categories: LP, ML, basal/fetal mix, repressive, and others (Table S3) (Figure 3D). Many factors were co-enriched in fetal and basal cells, suggesting the similarity of the transcriptional regulatory landscape of these two cell types. Consistent with previous findings (Dravis et al., 2015, 2018), members of the NF1 and SOX families are enriched in basal/fetal clusters, while ELF and FOX, JUN/FOS related factors are enriched in LP and ML clusters, respectively (Figure 3A and 3D).

We arranged these TFs into single-cell correlation networks to help visualize the similarity between the TFs, their corresponding clusters, and cell type annotations (Figure S7A). Specifically, ML- and basal/fetal-associated factors are at opposite ends of the network, indicating they are highly dissimilar. LP-specific factors are centrally positioned, suggesting an intermediate level of relatedness along the luminal-basal differentiation axis. Of note, although some transcriptional repressors such as SNAI2, ID3/4 and ZEB1 cluster with other luminal factors, these TFs likely function as basal factors by serving as repressors of luminal characteristics, as indicated by their chromatin accessibility profiles and previous reports (Phillips et al., 2014). Interestingly, in addition to the more well-known factors, we also identified TFs that have not typically been associated with mammary cell state. Examples of these include MAZ, SP2 and ZFP148 in fetal cells; EGR2/4 and CEBPB in basal cells; PPARG and MEIS1 in ML cells; EHF in LP cells; ZFP105 in all luminal cells; NFKB1 and RELA in all adult cells (Figure 3E and S7B). mRNA expression of some of these factors correlate well with their TF z-scores (Figure 3F), suggesting that these TFs may both be subject to cell-type specific regulation, and that they may also contribute to cell type determination. The combination of TF transcriptome data (Dravis et al., 2018; Giraddi et al., 2018), single-cell TF profiles, and chromatin accessibility data presented here provide a valuable resource for future characterization of candidate mammary cell state regulators. However, we note that members within the same TF family may have similar DNA binding motif profiles, and thus may not always be distinguishable, and that functional validation will be required for more precise assignment of regulatory roles.

### Mammary differentiation trajectory inferred by pseudotime ordering of single-cell TF profile

A ‘local’ dimensionality reduction method such as t-SNE enables cluster identification in single-cell data. However, t-SNE does not reveal cell state developmental trajectories that occur during tissue development. We thus used the Monocle 2 algorithm to organize single mammary cells into a ‘pseudotime’ trajectory based on their TF profiles (see Methods) (Qiu et al., 2017). Monocle 2 does not assume branch number and learns single-cell trajectory in a fully unsupervised manner. We again used our cell ID scores to identify the likely state of each branch and end point (Figure S8A). Our analysis reveals a continuum of developmental states between E18 fetal cells and post-natal luminal and basal cells, in which the LP state branches off halfway between the fetal and ML state. The closest cell state to the fetal is the basal, followed by LP and ML (Figure S8B). This is consistent with previous reports that LP and basal cells share transcriptional and epigenetic similarity to fetal cells (Dravis et al., 2018). Cells of intermediate state are also observed spanning between end states, demonstrating the levels of heterogeneity and potential plasticity within the mammary gland. Interestingly, we observed that some fetal cells are already entering the basal, intermediate or LP state, but rarely the ML state (Figure 4A), which is concordant with the cell ID score and TF z-score analyses. Superimposing TF z-score onto the pseudotime trajectory, we observed enrichment of state-specific factors in their associated cell type (Figure 4B). We also applied Monocle2 pseudotime ordering on our previously generated mammary gland scRNA-seq data (E16, E18, P4 and adult stages) (Giraddi et al., 2018). This resulted in a similar pseudotime trajectory (Figure S8C-E). Together, these observations validate our approach and show that the differentiation trajectory projected by this algorithm is consistent across different analytical approaches.

**Figure 4.**
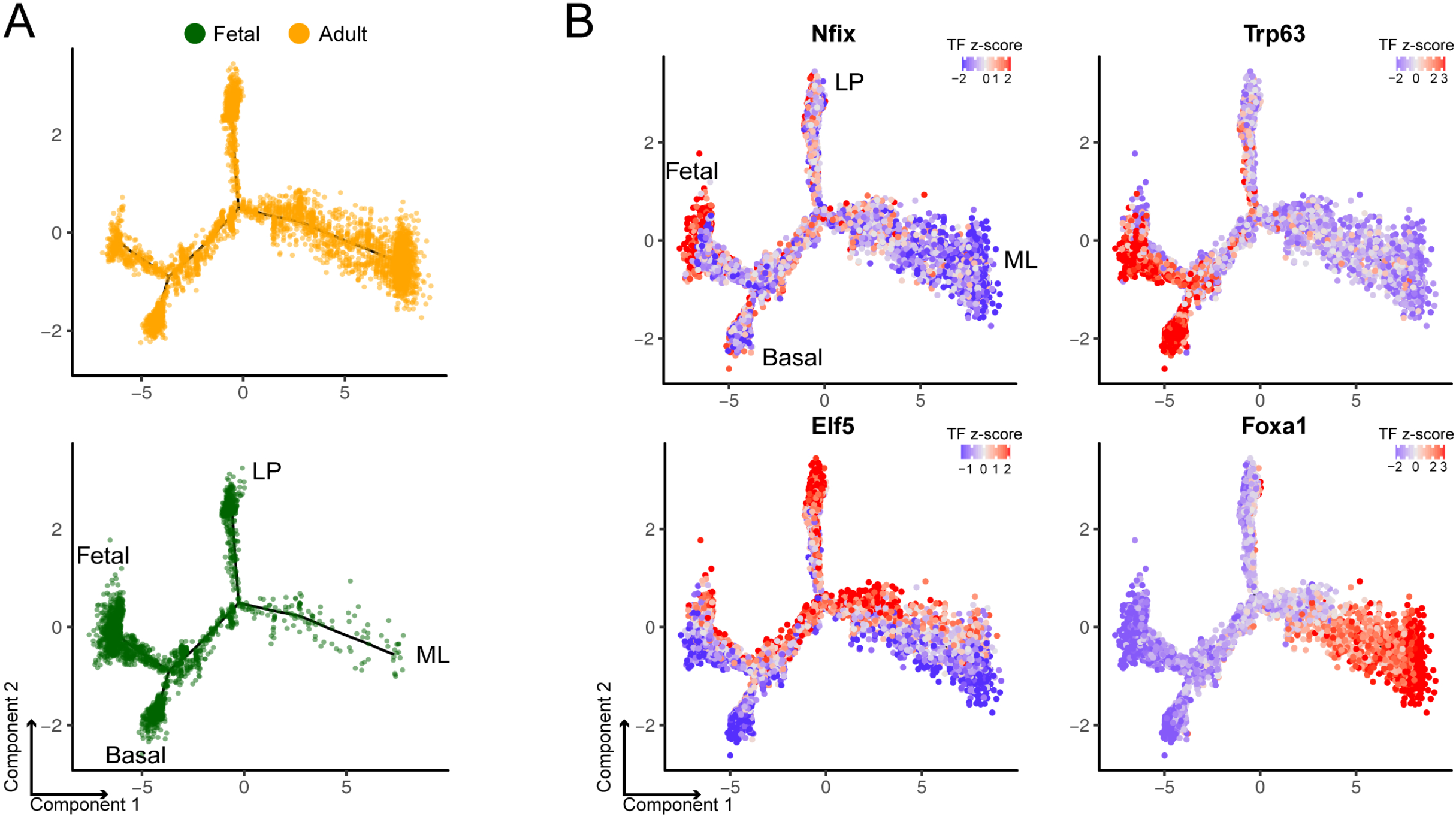
Pseudotime ordering of single-cell TF profile infers mammary differentiation trajectory. (A) Fetal (green) and adult (yellow) cells along the DDRTree pseudotime trajectory. Cell state associated with each branch is indicated. (B) Representative TF z-score profile overlaid on the pseudotime trajectory plot.

### Predicting cis-regulatory chromatin interactions in mammary cells

Chromatin accessibility changes at distal enhancers are generally more strongly correlated with cell-type than changes at gene promoters. Such changes can illuminate regulatory enhancers, suggest factors involved in altering gene expression to elicit phenotype, and indicate target genes of such regulation. Deciphering which of the many putative upstream elements comprise regulatory enhancers, and which of the potential targets they control are crucial for gaining a more precise understanding of the epigenetic and transcriptomic underpinnings of cell state regulation. However, the ‘nearest gene’ approach of mapping enhancer-promoter interactions is suboptimal because enhancers can be separated by significant distances from genes (Corces et al., 2018). Moreover, critical enhancer-promoter interactions may involve bypassing adjacent genes to reach the intended target.

We used the Cicero algorithm to identify putative enhancer-gene interactions relevant to cell state determination (Pliner et al., 2018). Cicero uses single-cell chromatin co-accessibility to develop a genome wide cis-regulatory map that is concordant with chromatin interaction data. Cicero identified 111,604 putative sites above the co-accessibility threshold of 0.2 in our snATAC-seq database. To validate the predicted co-accessible sites, we focused on the *Sox10* locus, as *Sox10* enhancers have been functionally characterized in the developing mouse embryo and in transgenic zebrafish (Antonellis et al., 2008; Betancur et al., 2010). We found that many of the enhancers that have strong putative co-accessibility with the *Sox10* promoter were also functionally verified as the main drivers of Sox10 expression (Figure 5A). These strong enhancers also overlap with the histone activation mark H3K27ac from published ChIP-seq data (Dravis et al., 2018) (Figure 5A), indicating that the Cicero predicted sites correlate with sites of known regulatory importance. In addition to the *Sox10* locus, we also identified putative co-accessible sites at important mammary cell state indicator genes. For example, we found a large enhancer cluster enriched with the activating H3K27ac mark connecting to both *Krt8* and *Krt18* genes (Figure 5B). This observation suggests the potential importance of this enhancer cluster in regulating luminal cell state. We also found putative distal sites that may be important in regulating genes involved in LP *(Kit)* and basal (*Krt5* and *Krt14*) state regulation (Figure S9). Collectively, these data indicate how chromatin accessibility mapping can serve as a valuable resource to predict factors relevant to mammary cell state determination.

**Figure 5:**
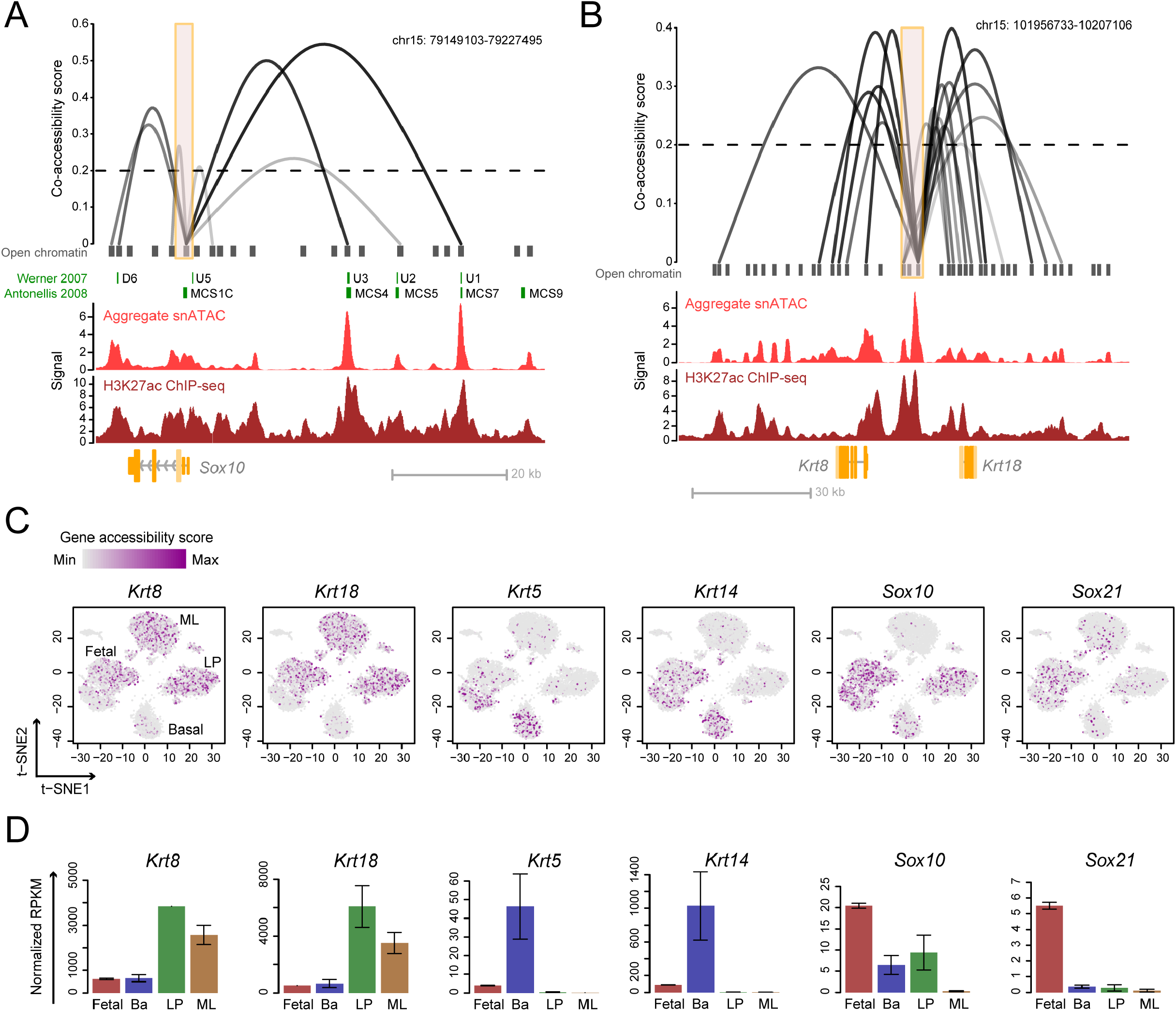
Putative cis-regulatory interactions and gene accessibility derived from snATAC-seq data. (A-B) Linkage plots of Cicero predicted cis-regulatory interactions at *Sox10* (A) and *Krt8/Krt18* (B) promoters (orange boxes indicate <±1.5 kb TSS). Previously characterized *Sox10* enhancers are shown in green. Signal tracks from aggregate snATAC-seq and H3K27ac ChIP-seq of fetal cells are shown. The height and opaqueness of the loop corresponds to the coaccessibility score between two linked elements. Dotted lines indicate co-accessibility cutoff. (C) Single-cell gene accessibility score of cell markers overlaid on the snATAC t-SNE plot. Cell cluster identities are annotated. (D) RNA-seq expression of genes from (C) in mammary cell types. Mean ± SEM (n = 2).

### Quantifying genome-wide gene accessibility at single-cell resolution

We used the inferred chromatin interaction map to profile the overall accessibility of each gene by quantifying the accessibility of all enhancers and their linked gene. This approach yielded a ‘gene accessibility score’ for 18,938 individual genes across 7,846 single cells. We filtered out genes that are not expressed in mammary cells based on RNA-seq, resulting in accessibility profiles for 11,327 genes (see Methods). Superimposing the gene accessibility score on the single-cell t-SNE plot shows enrichment of luminal markers *Krt8* and *Krt18* in both LP and ML, while basal markers *Krt5* and *Krt14* are enriched in basal cells (Figure 5C). In addition, the fetal associated genes *Sox10* and *Sox21* are most accessible in fetal cells (Figure 5C). In each of these cases, the chromatin accessibility score correlates well with mRNA expression (Figure 5D). These data validate the utility and specificity of the gene accessibility score as a resource to examine genome-wide gene accessibility. Importantly, we observed opposite enrichment of luminal (*Krt8/18*) and basal (*Krt5/14*) markers in the fetal sub-clusters that show characteristics of adult lineages (Figure 5C).

### Identification of mammary cell-type specific accessible genes

To identify genes that are specifically open or closed in each mammary cell type, we employed a machine learning strategy to find genes whose accessibility can predict cluster identity (see Methods). We used elastic net (Zou and Hastie, 2005), a penalized model that is suitable for high dimensional data learning but is more computationally efficient than random forest in processing the large number of genes presented here (Figure 6A and S10A). Our approach effectively identified top genes that are specifically open or closed for fetal, adult basal, LP and ML cells (Figure 6C, S10C and Table S4). The accessibility of these genes also correlates well with gene expression (Figure 6B and S10B), suggesting a connection between chromatin accessibility and cell-type specific gene expression pattern. Interestingly, many basal- or LP-specific genes also have increased levels of accessibility in fetal cells, again indicating the partial basal- and LP-specification of fetal cells and the epigenetic closeness between these cell types. Cluster 6 (Figure S3B) identified above was unusual in that it includes both fetal and adult cells but is distinct from the known epithelial cell types. We found that genes involved in ECM interactions and collagen formation are enriched in this cluster (Figure S11, Table S4 and S5). Given that this cluster also generally lacks accessibility at epithelial cytokeratins (Figure 5C), we infer that it represents stromal cell contamination or an unknown mesenchymal-like population. Further validation of this analytical method came from our identification of previously known cell type specific genes, such as *Sox11* in fetal, *Acta2* in basal, *Kit* in LP and *Foxa1* in ML (Figure 6D, left). We also identified new cell-type correlative genes, such as *Igf2bp3* in fetal, *Cxcl14* in basal, *Itga2* in LP and *Abcc8* in ML cells (Figure 6C, 6D and Table S4). Gene ontology (GO) analyses of these gene groups show that fetal cells are open with genes relating to extracellular matrix (ECM), transcriptional activity and tissue development; basal cells are open with genes of ECM, migration and motility; LP cells are open with genes of secretion, lipid and stimulus responses; ML cells are open with genes of signaling processes and mammary gland development, while closed with genes involved in cell motility, lipid response and ECM, all of which are terms associated with basal and LP identity (Figure 6E, S10D and Table S5).

**Figure 6.**
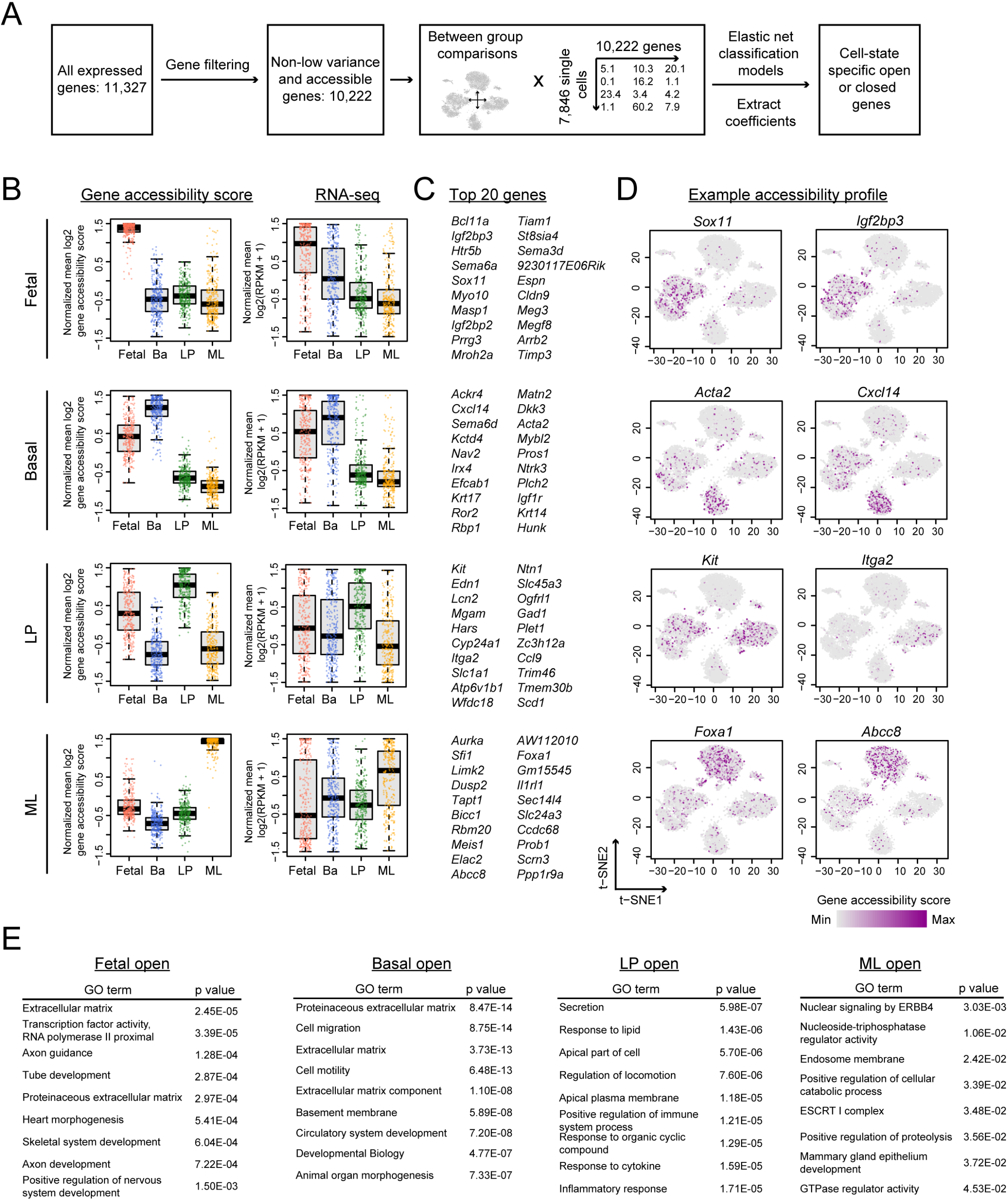
Identification and analysis of mammary cell-type specific accessible genes. (A) Computational framework to identify cell-type specific open or closed genes. (B) Boxplot of gene accessibility score and RNA-seq expression of top 300 accessible genes in fetal, basal, LP and ML cells. Each dot is one gene; thick horizontal middle line is the median; height of the box is the interquartile range (IQR); dotted vertical line is 1.5 x IQR. (C-D) Top 20 genes (C) and their representative accessibility profile (D) from (B). (E) Gene ontology (GO) analysis of top 300 accessible genes from each cell type. P values are Bonferroni corrected for multiple testing.

To identify differentially accessible genes between the fetal basal-like and LP-like cells, we used random forest instead of elastic net because of the relatively smaller number of cells in these two fetal clusters. Using a stringent cutoff based on checking the ranked profile on t-SNE plot, we selected the top 65 most important genes as signature genes of fetal basal-like and fetal LP-like cells (Table S4). Our approach successfully identified previously known basal lineage-associated genes, such as *Trp63, Krt5,* and *Acta2,* and Luminal lineage-associated genes, such as *Ehf, Krt8, Krt18,* and *Krt19.* By overlaying the enrichment score of the fetal basal-like signatures and fetal LP-like signatures on our previously generated scRNA-seq data of E16, E18, P4 and adult mammary cells, we found that, although E18 cells remains in one tight cluster, they start to show signs of basal and luminal lineage bifurcation (Figure S12B). In contrast, fetal E16 cells remain in one single population with no significant enrichment of these basal-like and LP-like signatures. This validates our snATAC-seq analytical approaches, and more importantly, further indicates that snATAC-seq better differentiates cell states during development. In addition, we found cell surface genes that exhibited differential accessibility between basal- and LP-like fetal cells (Figure S12C), and that were also differentially transcribed between basal- and LP-like fetal E18 cells (Figure S12D). They may serve as candidate markers for separating the fetal subpopulations for future functional studies.

### An online resource for visualization of mammary snATAC-seq data

We developed and launched a web application for our snATAC-seq data to serve as a scientific resource for further investigation of genes and regulatory mechanisms involved in mammary differentiation, and to facilitate data visualization, (https://wahl-lab-salk.shinyapps.io/Mammary_snATAC/). Features such as cell ID score, TF z-score and gene accessibility score (11,327 expressed genes) across single mammary cells can be plotted with the t-SNE or pseudotime plots seen throughout this report. We have also included the published bulk RNA-seq data for easy comparison between chromatin accessibility and gene expression. Additionally, to facilitate the visualization of our cicero inferred putative promoter-enhancer interactions, we have generated a *.longrange* format connections file containing all high quality connections mapping to ±1.5kb TSS (Supplement file1, see Methods for instructions of visualization).

## Discussion

Our snATAC-seq computational framework can be scaled for larger single-cell epigenetic studies, but here we focused on two stages of mammary gland development (comprising late embryogenesis and the adult gland) and developed a reference epigenome database based on thousands of embryonic and adult virgin cell types. Analysis of the E18 embryonic cells enabled us to ascertain whether pre-natal mammary cells retain bipotent features, as evident in previous lineage tracing studies, or are lineage-specified before birth, as suggested by recent lineage tracing reports. Interestingly, our snATAC-seq analysis reveals that E18 fMaSCs contain luminal-oriented and basal-oriented fetal cells, with the fetal-like LPs and fetal-like basal cells being equidistant from the bulk fetal ATAC principal component coordinate. Taken together with the RNA-seq data, we interpret this finding to suggest that most of the cells at this stage are un- or weakly-committed, but already biased towards either a luminal or basal fate. Monocle 2 analysis of the snATAC-seq data is consistent with this interpretation. The pseudotime trajectory shows two clear branches: a basal branch that generates adult basal cells, and a luminal branch that generates an intermediate that coincides with the luminal progenitor. Mature luminal cells then descend from the luminal progenitor. This pseudotime analysis is not consistent with a prior suggestion of a basal intermediate for luminal progenitors (Pal et al., 2017), but is consistent with our and other single-cell RNA-seq analyses (Bach et al., 2017; Giraddi et al., 2018). It will be important to now use snATAC-seq to assess earlier embryonic stages to determine when the first evidence of luminal and basal orientations occur, and to deduce the cues leading to such bifurcation. Analyses of the epigenomes of cells undergoing pubertal expansion (P21-P35), hormone-induced changes in the adult during estrus cycling, pregnancy-associated changes such as development of the lactation-competent gland and involution, and the different subtypes of breast cancer should also be informative concerning how cellular and epigenetic heterogeneity correlate, and to infer the underlying regulators, which can then be functionally evaluated using genetic and other strategies.

The snATAC-seq pipeline we used enabled epigenetic profiling of thousands of adult mammary cells, which produced an agnostic assessment of the heterogeneity of the basal and luminal cell lineages. Studies have consistently shown that the basal mammary cells transplant to form full and functional outgrowths with an efficiency of ~1-2%. Luminal cells largely lack this capacity. One interpretation of this observation is that basal cells are heterogeneous and contain a minority subset correlating to a bipotential progenitor that is not revealed by lineage tracing. We therefore interrogated our large snATAC-seq database of adult basal cells to try to identify either a 1-2% sub-population that may possess these characteristics, or the predicted stem-enriched populations obtained by sorting for markers such as Lgr5, Procr, and Tspan8 (Fu et al., 2017; Plaks et al., 2013; Wang et al., 2015). Our chromatin profiling data, however, show that basal cells comprise a single cluster. Thus, while there is heterogeneous expression of these markers within the basal cell fraction, our data suggest that such expression does not correspond to significant differences in net chromatin accessibility or the existence within the basal population of a subset with the characteristics of the highly plastic fMaSC population.

Our research group and others have previously performed single-cell transcriptomic profiling on mammary gland cells at similar stages of development (in the embryo and adult). In agreement with the chromatin profiling data described here, these analyses also found clear distinction between the three-separate basal, LP, and ML lineages of the adult mammary gland. Interestingly, our scRNA-seq analysis of embryonic mammary cells revealed that, despite significant transcriptional heterogeneity, E18 mammary cells presented as a single net population (Giraddi et al., 2018). By evaluating the enrichment of fetal, basal- and LP-like signature genes inferred from the Cicero gene accessibility scores, we found that, although E18 cells remain in one tight cluster, they start to show signs of basal and luminal lineage bifurcation. By contrast, E16 mammary cells remain in one single population with no enrichment of adult basal- and LP-like signatures. It is possible that the epigenetic changes that are occurring at E18 have yet to be fully reflected at the transcriptomic level due to the lag time between generating an epigenetic change and the consequent change in transcription. Other factors that may complicate deduction of cell state inferred by RNA include the stability of historically produced mRNAs, and the production of key regulatory transcription factors below the level of detection. Consistent with the lag between change in chromatin and transcriptional readout, we note that mammary cells from postnatal day 4 mice show clear separation of basal and luminal populations (Figure S12B). These results together indicate that snATAC-seq is particularly valuable in delineating developmental cell states and differentiation programs.

In our previous analyses of embryonic and adult mammary cells using bulk transcriptomic and epigenetic profiling (Dravis et al., 2018), we found that chromatin profiling indicated that LP were the adult cell type that most closely resembled the chromatin features associated with fMaSCs. This was of interest because LPs have consistently been suggested as the cell of origin for the aggressive basal-like breast cancer subtype, which we have previously shown possess transcriptomic similarities to fMaSCs (Lim et al., 2009; Molyneux et al., 2010). The close relationship between fMaSCs and LP is consistent with the single-cell chromatin analyses presented here, as we find that adult LPs profile more closely to LP-like fMaSCs and bulk fMaSCs than do adult basal cells by PCA. We also find that the majority of E18 cells have adopted LP-associated chromatin features. Interestingly, snATAC-seq also revealed that LP uniquely exhibit a number of features associated with a basal cell identity, and such features are not apparent in the ML population. Thus, LP may inherently possess fMaSC-like and mixed-lineage characteristics associated with cell state plasticity, which may explain their association with being a preferred cell of origin of basal-like breast cancers in several mouse models, and of their ability to acquire expanded cell state plasticity in response to activation of several different oncogenes (Koren et al., 2015; Van Keymeulen et al., 2015).

The single-cell epigenetic data presented constitute a technical resource for deconstructing the cell state heterogeneity of a developing tissue and for identifying putative regulatory elements and transcription factors involved in cell type specification. Our chromatin profiling of individual mammary cells at embryonic and adult developmental stages, and accompanying analyses that predict transcription factor activity and gene accessibility in relation to distinct mammary cell states, provide valuable resources to discover and validate cell state regulators. The links between mammary development and breast cancer suggest that this resource, which we have made available as a web-based app to facilitate accession of, will have significant utility in target discovery for breast cancers.

## Methods

### Mammary cell isolation

Normal mammary epithelial cells were isolated based on previously described procedures. All cells were isolated from CD1 mice. E18 mammary rudiments (all rudiments), and 8 weeks old adult mammary glands (#4 gland only) were dissected and pooled from multiple animals into dissection media (Epicult-B Basal medium (Stem Cell Technologies #05610) supplemented with 5% FBS, pen/strep, fungizone, hydrocortisone, collagenase and hyaluronidase), and agitated with shaking for 1.5 hours for the fetal tissue, and 3 hours for the adult tissue at 37C. Erthyrocytes were removed with ammonium chloride exposure for 4 minutes on ice, followed by cell trituration with dispase and DNase. Final suspensions were passed through a 40 um filter to remove cell aggregates, and stored in Hank’s Balanced Salt Solution with 2% FBS for immunostaining and flow cytometry sorting. Cells were then stained with EpCAM-AF647 (BioLegend #AB_1134101) and lineage markers (Biotin-Ter-119 (BD #553672), Biotin-CD45 (BD #553078) and Biotin-CD31 (BD #553371)), with mouse Fc Block (BD #553141) on ice for 15 minutes, followed with lineage markers conjugation with SA-APC-Cy7 (BioLegend #405208). For cell sorting, DAPI+ and Lin+ cells were excluded, and total mammary epithelial populations were flow sorted as EpCAM+, using a BD InFlux cell sorter. Immediately after sorting, cells were resuspended in 10% DMSO, 20% FBS in DMEM and frozen at −80. To obtain enough cell number (>200k per sample) for the snATAC-seq protocol, each adult biological replicate was pooled from sorted cells of three adult mice, while each fetal biological replicate was pooled from sorted cells of approximately 80 fetuses.

### Combinatorial barcoding assisted single-nuclei ATAC-seq

Combinatorial ATAC-seq was performed as described previously with modifications (Cusanovich et al., 2015; Preissl et al., 2018). For the fetal samples, mammary epithelial cells from roughly 65 E18 embryos were pooled for each biological replicates. For the adult samples, cells from 3 8-weeks old mice were pooled per biological replicate. For each sample two biological replicates were processed. Cells were thawed and pelleted in a swinging bucket centrifuge (500 x g, 5 min, 4°C; 5920R, Eppendorf). Cell pellets were resuspended in 250 μl nuclei permeabilization buffer (10 mM Tris-HCl pH 7.4 (Sigma), 10 mM NaCl, 3 mM MgCl2, 0.1% GEPAL-CA630 (Sigma), 0.1% Tween-20, and 0.01% Digitonin (Promega, G9441), cOmplete (Roche) in water) (Corces et al., 2017), incubated for 5 min at 4 °C with rotation and pelleted again (500 x g, 5 min, 4°C; 5920R, Eppendorf). Nuclei were resuspended in 500 μL high salt tagmentation buffer (36.3 mM Tris-acetate (pH = 7.8), 72.6 mM potassium-acetate, 11 mM Mg-acetate, 17.6% DMF) and counted using a hemocytometer. Concentration was adjusted to 2000 nuclei/9 μl, and 2,000 nuclei were dispensed into each well of a 96-well plate – 32 wells for each of the adult replicates and 16 wells for the fetal replicates, respectively. All downstream pipetting steps were performed on a Biomek i7 Automated Workstation (Beckman Coulter). For tagmentation, 1 μL barcoded Tn5 was added (Picelli et al., 2014), mixed and incubated for 60 min at 37 °C with shaking (500 rpm). To inhibit the Tn5 reaction, 10 μL of 40mM EDTA were added to each well and the plate was incubated at 37 °C for 15 min with shaking (500 rpm). Next, 5 μL 5 x sort buffer (5 % BSA, 5 mM EDTA in PBS) were added. All wells were combined and filtered using a 30-μm CellTric (Sysmex) into a FACS tube and stained with 3 μM Draq7 (Cell Signaling). Using a SH800 (Sony), 20 nuclei were sorted per well into eight 96-well plates (total of 768 wells) containing 10.5 μL EB (25 pmol primer i7, 25 pmol primer i5, 200 ng BSA (Sigma)). After addition of 1 μL 0.2% SDS, samples were incubated at 55 °C for 7 min with shaking (500 rpm). We added 1 μL 12.5% Triton-X to each well to quench the SDS and 12.5 μL NEBNext High-Fidelity 2× PCR Master Mix (NEB). Samples were PCR-amplified for 12 cycles (72 °C 5 min, 98 °C 30 s, (98 °C 10 s, 63 °C 30 s, 72 °C 60 s) × 11, held at 72 °C). After PCR, all wells were combined. Libraries were purified according to the MinElute PCR Purification Kit manual (Qiagen) and size selection was performed with SPRI Beads (Beckmann Coulter, 0.55x and 1.5x). Libraries were quantified using a Qubit fluorimeter (Life technologies) and the nucleosomal pattern was verified using a Tapestation (High Sensitivity D1000, Agilent). The library was sequenced on a HiSeq2500 sequencer (Illumina) using custom sequencing primers, 25% spike-in library and following read lengths: 50 + 43 + 37 + 50 (Read1 + Index1 + Index2 + Read2). The library was sequenced twice to obtain enough reads depth.

### Single-nucleus ATAC-seq data processing and cluster analysis

All bioinformatic analyses were performed with bash script and R. Fastq files from the two sequencing runs were merged. Pair-end sequencing reads were trimmed with sickle (https://github.com/najoshi/sickle) to remove low quality base pairs, and mapped to mm10 reference genome with bowtie2 with the following parameters: bowtie2 -p 16 -t -X 2000 --no-mixed --no-discordant (Langmead et al., 2009). Data processing, which includes low quality reads filtering (MAPQ < 30), removal of duplicates and mitochondrial reads, adjusting for Tn5 insertion and separation of single-cell reads, were performed with the snATAC-seq pipeline as previously described (Preissl et al., 2018) with the following parameters: snATAC pre -t 16 -m 30 -f 2000 -e 75. Open chromatin regions were determined by calling peaks for each replicate using macs2 with these parameters: macs2 callpeak --nolambda --nomodel --shift -100 --extsize 200 --keep-dup all -- bdg --SPMR -q 5e-2 (Zhang et al., 2008). Peaks from each replicate were merged with bedtools into non-overlapping regions (Quinlan and Hall, 2010), resized to 1 kb and separated into promoter proximal (< ±1.5 kb TSS) and distal (>= ±1.5 kb TSS) regions. For cell selection, we filtered out cells that did not pass our quality threshold: reads per cell >= 2000 and reads in peak ratio >= 0.2. Data statistics for each biological replicate is shown in Table S1. The aggregate snATAC profile was plotted with UCSC genome browser (https://genome.ucsc.edu), and the single-cell reads profile was plotted with the R package Sushi. Average signal profile was plotted with deepTools (Ramírez et al., 2014).

To reveal single-cell clusters with proper data normalization, we adapted a previously described workflow to process the snATAC-seq data (Cusanovich et al., 2015). First, we calculated a binary matrix of cell versus open chromatin regions using the snATAC pipeline tool snATAC bmat, with 1 indicating >= 1 reads and 0 indicating < 1 reads within a specific region. We next performed latent semantic indexing and singular value decomposition, keeping only the top 50 dimensions. Of note, since we did not observe strong correlation between PC1 and read depth as described previously (Figure S2A), we kept all 50 dimensions for further analysis. Next, we performed dimension reduction with t-distributed stochastic neighbor embedding (t-SNE) (Rtsne package in R; parameters: dims = 2, perplexity = 30, max_iter = 1000, theta = 0.5, pca = FALSE, exaggeration_factor = 12), and uniform manifold approximation and projection (UMAP) (umap package in R; parameters: n_components = 2, n_neighbors = 15, random_state = 524). Unbiased cluster identification was performed with density peak clustering using the R package densityClust, with rho = 50 and delta = 3.5 (Rodriguez and Laio, 2014).

Previous reporting shows that our snATAC-seq procedure may generate barcode collision (Preissl et al., 2018), meaning that reads from two different cells possess the same barcode. As library collision may confound cell type identification, we excluded cell clusters whose library complexity (defined as numbers of unique reads per cell) are significantly higher than expected. To do so, we sampled the population with cell number corresponding to each cluster for 10,000 times, and plotted this background library complexity profile with 99% confidence intervals against the actual profile from each cell cluster. Clusters that have significantly higher library complexity distribution than the background were considered as barcode collision cells, and were not further analyzed.

To annotate cluster identity, we calculated single-cell normalized accessibility at previously defined cell-type specific regions: uniquely accessible regions (UARs) and uniquely repressed regions (URRs) (Dravis et al., 2018). We first resized the UARs/URRs to 1 kb based on peak center so that downstream calculation will not be biased by peak size. We next generated a binary matrix of cells versus UARs/URRs using snATAC pipeline described above, and calculated the reads depth normalized ‘cell ID score’ defined as: (sum of UAR counts / reads per cell) – (sum of URR counts / reads per cell). This cell ID score was calculated for each single-cell and for each cell type UARs/URRs, including those of fetal, basal, LP and ML cells. To reduce outlier effect and better reveal the intermediate range, the cell ID score was capped at upper and lower 10% quantile and plotted.

### Transcription factor dynamics, clustering and network analysis

TF dynamics were inferred with chromVar package as previously described (Schep et al., 2017). Open chromatin regions called from aggregate profile described above was used to calculate read counts at open chromatin. mm10 reference genome was used for correcting GC bias, and chromVar curated motif V2 (mouse_pwms_v2) was used for generating TF z-score. A total of 788 TFs were used for downstream analysis. TF variability was calculated as the variance of TF z-score across all the single cells, and it is compared against a ‘expected variability’ calculated by the mean of the same TF z-score profile permutated for 1000 times. TF z-score score was capped at upper and lower 10% quantile when plotted to reduce outlier effect and better reveal the dynamic range.

We employed a random forest classification framework to find TFs whose TF z-scores predict major cell types. We used random forest because it is highly accurate, robust to noise and is suitable for high dimension data modeling. When applied with permutation importance (rather than Gini importance), random forest variable analysis is relatively resistant to bias caused by predictor co-linearity (https://explained.ai/rf-importance/index.html#6.1). We used the R package Caret (http://topepo.github.io/caret/index.html) to individually tune our models and extract important variables, with these parameters: mtry = 5-200, ntree = 500, importance = TRUE, metric = ROC. The predictors are the 788 TFs, with read depth per cell as a covariate. The class comparisons for the model are: fetal vs. all adults, basal vs. all other adults, LP vs. all other adults, ML vs. all other adults, and basal only vs. ML only. To improve accuracy, cells corresponding to the stromal contamination (cluster 6 in Figure S3B) and barcode collision doublets (cluster 10 and 14 in Figure S3B) are excluded from random forest analysis. We excluded fetal cells in the adult classification model as fetal cells show characteristics of adult cell types, and thus may confound our model. The top TFs from each model were further checked by visualization to ensure the effectiveness of the variable analysis.

To cluster TFs based on their single-cell profile, we pooled the top TFs from the random forest variable analysis to generate a list of cell-type predictive TFs, and then performed clustering on these key TFs. The pooling was done with the top 50 TFs from each classification model, except for the ‘basal only vs. ML only’ model, which we selected the top 100 TFs. The pooling results in 148 TFs. We next performed K-medoid consensus clustering with Pearson correlation on these 148 TFs based on their chromVar TF z-score, using the R package ConsensusClusterPlus (Wilkerson and Hayes, 2010), with these parameters: maxK = 15, reps = 1000, pItem = 0.8 (80% of TFs sampled), pFeature = 0.5 (50% of cells sampled), distance = pearson, clusterAlg = pam, seed = 2018. We used cophenetic coefficient to determine the optimal cluster number. The median TF z-scores for each TF cluster in fetal-Basal-like, fetal-LP-like, fetal-ML like, basal, LP and ML cells were plotted to check the enrichment profile.

We constructed a weighted pearson correlation network with the TF z-score across the single cells, using the R package igraph (Csárdi and Nepusz, 2006). We removed weak edges that have correlation < 0.2, and used the Fruchterman Reingold algorithm to construct the network.

### scRNA-seq data processing

10x scRNA-seq data was download from SRA with accession number GSE111113. Fastq files were processed with cellranger v3.0.1 using default parameters. Then the filtered feature barcode-count matrix of each stage was read into Seurat object for cell quality check and filtering. Individual Seurat object of each stage were combined into a single Seurat object by using the MergeSeurat (Butler et al., 2018) function with do.normalize = FALSE. Cell selection in Seurat was performed as described previously by selecting the percent of mitochondria (low.thresholds = -Inf, high.thresholds = 0.075), number of UMI per cell (low.thresholds = 200, high.thresholds = 50000), and number of genes detected (low.thresholds = 200, high.thresholds = 7000). This resulted in 691 E16, 1455 E18, 1377 P4, and 2250 adult high quality single cells. Then the expression matrix was read depth normalized and log transformed with the NormalizeData function of Seurat package (Butler et al., 2018). Dimension reduction and visualization was done in R using the UMAP package with default parameters. Top 1000 most variable genes across all the single cells were used. Enrichment of snATAC gene accessibility derived fetal Ba-like and LP-like signature genes in each individual cell was evaluated by using the AUCell R package (Aibar et al., 2017) with the default settings. Then the fetal Ba-like and LP-like signature enrichment scores were overlaid on the UMAP dimension 1 and dimension 2 plot.

### Pseudotime analysis

We used Monocle2 with the TF z-score profile for pseudotime ordering of single cells as described previously (Qiu et al., 2017). The top 200 most variable TFs across all the single cells were used for ordering cells based on the ranked TF z-score variability (Figure S6A). Dimension reduction and trajectory learning were performed with these parameters: max_components = 2, method = DDRTree, norm_method = none; the cell ordering was performed with the default settings. For scRNA-seq pseudotime trajectory, the filtered gene count matrix was imported from the processed expression matrix in the Seurat object with the importCDS function of monocle2 package (Qiu et al., 2017). Top 1000 most differentially expressed genes across 12 clusters were selected for ordering cells in pseudotime. Cluster numbers were selected based on the tSNE plots. Cells were reduced into 3 DDRTree dimensions (norm_method = "log", reduction_method = "DDRTree", max_components = 3) and then ordered in pseudotime with default parameters. Branch of E16 cells was set as the root.

### Prediction of cis-regulatory interaction and gene accessibility analysis

Cicero analysis was performed as described previously (Pliner et al., 2018). We input Cicero with binarized chromatin accessibility at all promoter and distal regions as defined above. We used the t-SNE coordinates calculated above for dimension reduction and nearest neighbor calculation, with mouse mm10 as the reference genome. In total, we revealed 3,622,246 chromatin interactions with Cicero co-accessibility score > 0. The R package Gviz was used for plotting cis-regulatory interaction maps and genomic tracks (Hahne and Ivanek, 2016).

To calculate genome-wide gene accessibility score, we used the Cicero function ‘build_gene_activity_matrix’, with co-accessibility cutoff = 0.2. Instead of only using promoter regions, we used UTRs and exons to annotate genes and then calculated gene activity scores for each single cell. Comparing to only using promoters, using UTRs and exons improves the results in several aspects: 1) gene transcribed from alternative promoters are captured; 2) it gives better signal coverage across single cells while not compromising cluster-specificity (examples are shown in Figure S13A). 3) it covers more genes (11327 vs 11063) and gene accessibility score of these additional genes are largely concordant with RNA-seq data (examples from the top 20 open genes are shown in Figure S13B,C). We filtered out nonexpressed genes based on published bulk RNA-seq data (Dravis et al., 2018), resulting in 11,327 genes with accessibility score. Non-expressed genes were defined as those that have a RPKM value < 3 in all normal and tumorigenic mammary cells. For visualization, gene accessibility scores were capped at upper and lower 5% quantiles to reduce outlier effect and better reveal the intermediate range.

To identify cell-type specific open or closed genes, we applied an elastic net classification framework on the gene accessibility score. We used elastic net instead of random forest as described above, because the latter is inefficient with large numbers of predictors. Although elastic net is generally less accurate and flexible than random forest, the penalization function of this algorithm allows effective variable importance analysis even when co-linearity is present. We first filtered out low variance and low accessibility genes of which their accessibility scores variance and sum across the single cells are at the bottom 5%, which results in 10,222 genes being kept. We next used the R package Caret to perform elastic net tuning with these parameters: method = glmnet, preProc = c("center", "scale"), tuneLength = 3, metric = ROC. The predictors are the 10,222 genes, with 7846 single cells as samples. The class comparisons for the model are: fetal vs. all adults, basal vs. all other adults, LP vs. all other adults, ML vs. all other adults, and basal only vs. ML only. The top open and closed genes in each cell type are defined as those having the highest and lowest elastic net coefficient, respectively. The top genes were checked by visualization to ensure the effectiveness of the variable analysis. Gene ontology analysis was performed with ClueGO network analysis under Cytoscape (Bindea et al., 2009; Shannon et al., 2003), with these term libraries: GO molecular function, GO cellular component, GO biological process, KEGG, Reactome and Wiki pathway.

To get the fetal Ba-like and LP-like accessible signature genes, we used random forest instead of elastic net due to the smaller number of fetal cells. To avoid potential outliers’ effect on data centering and scaling, we capped the Cicero gene accessibility scores at top and bottom 1% before running random forest. We used the R package Caret to tune the model and extract important variables, with these parameters: mtry = 5-200, ntree = 500, importance = TRUE, metric = ROC. Tuned best mtry=200, AUC=0.9151 with 95%CI: 0.9037-0.926 (DeLong). Accessibility profiles of top variables were visually checked on t-SNE plot and the top 65 genes were selected as the most important signature genes to distinguish fetal Ba-like vs LP-like cells. Stringent cutoff was applied to reduce false positives. Each of these 65 genes were assigned to Ba- or LP-like class according to its mean expression in these two clusters.

To calculate the ‘basal-to-luminal score’, top 300 basal and ML specific genes from the ‘basal only vs. ML only’ model are extracted. For each cell, the score is calculated as: (sum of accessibility score at basal specific genes) / (sum of accessibility score at luminal specific genes). For visualization on the t-SNE plot, the scores were capped at upper and lower 10% quantiles.

To perform principle comportment analysis (PCA) with bulk ATAC-seq and aggregate snATAC-seq profile, we used deepTools to calculate ATAC signal enrichment of each sample/cell type at all the UARs and URRs. For normalization, the sample-wise signals were centered and scaled, while the region-wise signals were centered, and then subjected to PCA with the R function ‘svd’. The 3D PCA plot was plotted with the R package Rgl (https://cran.r-project.org/web/packages/rgl/vignettes/rgl.html).

### Mammary snATAC-seq web application

The snATAC-seq web application is written with R Shiny and served on the Shiny server (https://www.rstudio.com/products/shiny/).

### Cicero connections visualization on WashU Epigenome Browser

All high quality cicero connections (accessibility score >= 0.2) that map to peaks that overlap with protein-coding gene promoter regions (±1.5kb TSS) were converted into *.longrange* file format, bgziped, and tubix indexed for convenient visualization in WashU Epigenome Browser. The accessibility score of each single cell generated by cicero was scaled by 100x for easy plotting. To visualize the cicero connection in WashU Epigenome Browser (http://epigenomegateway.wustl.edu/browser), both the *.gz* file and the *.gz.tbi* files in supplement file1 can be uploaded as a local track and visualized in “ARC” track display mode.

### Data and software availability

Raw and processed data have been deposited to NCBI Gene Expression Omnibus with the accession number GSE125523. Previous published 10x scRNA-seq data was download from SRA with accession number GSE111113.The basic scripts for snATAC-seq analysis can be accessed here: https://github.com/jaychung10010/Mammary_snATAC-seq

## Acknowledgements

We thank Cynthia Ramos and Luke Wang for laboratory management, Grace Lee and Annie Odelson for laboratory assistance, Conor Fitzpatrick and Caz O’Connor at the Salk Flow Cytometry Core, Max Shokirev at the Salk Bioinformatics Core, Nasun Hah at the Salk Next Generation Sequencing Core, and Nikki Lytle and Eugene Ke for critical evaluation of the manuscript. Work in the laboratory of G.M.W. was supported, in part, by the Cancer Center Core Grant (5 P30CA014195), National Institutes of Health/National Cancer Institute (R35 CA197687), the Susan G. Komen Foundation (SAC110036), the Leona M. and Harry B. Helmsley Charitable Trust (2012-PG-MED002), the Breast Cancer Research Foundation (BCRF), The Freeberg Foundation, The Copley Foundation, and The William H. Isacoff MD Research Foundation for Gastrointestinal Cancer. C.D. was supported by CA174430. B.R. was supported by the Ludwig Institute for Cancer Research.

## Author Contributions

C-Y.C., C.D., and G.M.W. designed the experiments. C-Y.C., G.L., S.P. and X.H. performed snATAC-seq. C-Y.C. and Z.M. performed bioinformatic analyses with help from O.P. All authors contributed to the interpretation and presentation of experimental results. C-Y.C., C.D., Z.M. and G.M.W. wrote the manuscript. B.R. and G.M.W. provided funding.

